# Metabolic Signatures of Pulmonary Embolism in COVID-19: Insights from Longitudinal Intensive Care Unit Profiles

**DOI:** 10.1101/2025.09.12.675971

**Authors:** Merys Valdez, Malou Janssen, Zhengzheng Zhang, Lieke Lamont, Madhulika Singh, Wei Yang, Charlotte Nijgh van Kooij, Jelte J.B. Geerlings, Ahmed Ali, Naama Karu, Yolanda de Rijke, Amy Harms, Alida Kindt, Diederik Gommers, Henrik Endeman, Thomas Hankemeier

## Abstract

**Background:** Pulmonary embolism is a severe complication of COVID-19 infection, associated with a hypercoagulable state and heightened risk of blood clots. As SARS-CoV-2 has become endemic, understanding pulmonary embolism’s metabolic effects in COVID-19 patients is warranted. This study investigated the longitudinal metabolic profiles of 66 Intensive Care Unit-admitted COVID-19 patients at Erasmus Medical Center to identify metabolites and mechanisms associated with pulmonary embolism.

**Method:** A total of 1209 metabolic species were measured, including amines and lipids. Metabolic changes were analysed across four timeframes: i) general analysis of pulmonary embolism, ii) 72 hours prior to pulmonary embolism, iii) 48 hours prior and the day of pulmonary embolism, and iv) the day of and 48 hours post-pulmonary embolism.

**Results:** The general analysis revealed significant upregulation of amines, triglycerides, phosphatidylethanolamines, ether-linked phosphatidylethanolamines, and eicosanoids in patients who developed a pulmonary embolism. Phosphatidylethanolamines containing the 20:3 fatty acid side chain were notably elevated. Minimal metabolic dysregulation was observed 72 hours before pulmonary embolism, with subtle increases in lysophosphatidylcholines and lysophosphatidylethanolamines. In contrast, there was a strong metabolic response during and post-pulmonary embolism, phosphatidylethanolamines (47%), ether-linked phosphatidylethanolamines(96%) and sphingosines(40%).

**Conclusion:** These findings underscore the critical role of lipid metabolism in pulmonary embolism, particularly triglycerides and specific lipid species. The limited metabolic perturbations before pulmonary embolism suggest early prediction challenges, emphasising the need for further research into temporal metabolic changes and their clinical applications.

## 1. INTRODUCTION

The COVID-19 pandemic has challenged healthcare systems worldwide, and researchers are still trying to understand the multifaceted aspects of SARS-CoV-2 infection. Beyond its initial characterisation as primarily a respiratory illness,[1] COVID-19 has revealed its complex and systemic nature, affecting various organs and systems throughout the human body[2,3] One life-threatening complication of severe COVID-19 infection is pulmonary embolism [4].This condition arises when a thrombus—most commonly in the lower extremities—is transported to the pulmonary arteries and obstructs the arterial lumen. In some cases, thrombi may also form directly within the lungs [5–7]. Pulmonary embolism is controlled by a complex coagulation cascade that includes various factors and can be initiated by plasma hypercoagulability, alterations in blood flow and endothelial cell dysfunction [8].

In the initial stages of the SARS-CoV-2 pandemic, clinical investigators in Wuhan, China, reported a significantly heightened risk of thrombosis in COVID-19 patients [9]. Additionally, the elevated risk of thrombosis persists for as long as a year after COVID-19 diagnosis [10]. This observation was later corroborated by additional studies, supporting the notion that COVID-19-associated coagulopathy (CAC) plays a role in the increased morbidity and mortality among those infected with SARS-CoV-2 [11–18]. CAC can present as both microthrombi and macrothrombi, damaging several organs, such as the lungs, heart, brain and kidneys [18,19].

Computed tomography pulmonary angiography (CTPA) has become the first-line imaging modality for diagnosing pulmonary embolism [20,21]. It offers high sensitivity and specificity, particularly with multi-detector CT technology, which improves visualisation of peripheral pulmonary arteries and small emboli [22]. With its high negative predictive value, CTPA enables clinicians to confidently withholdanticoagulation treatment in patients with negative findings [21].

However, CTPA is associated with radiation exposure, contrast-related complications and its cost can vary widely depending on the healthcare setting, payer mix, and downstream resource utilization [23,24]. To mitigate these risks, a standardised diagnostic algorithm incorporating clinical decision rules and D-dimer testing can reduce unnecessary CTPA use by 20-30% [23]. Such CT angiographies are performed when clinical suspicion of pulmonary embolism is high or when D-dimer levels are increased above prior set limits [25,26]. While elevated D-dimer levels are associated with pulmonary embolism, they have a low positive predictive value and cannot confirm its presence. D-dimer testing alone is insufficient to confirm pulmonary embolism. Instead, the potential benefit lies in a negative test result’s ability to rule out the diagnosis [26].

The negative predictive value of D-dimer depends on assay sensitivity and inversely correlates with venous thromboembolism prevalence [26]. D-dimer is often elevated in hospitalised patients and those with severe infections or inflammation [27]. Thus, it has low-reliability for ruling out pulmonary embolism in high-risk patients, even with sensitive assays. In COVID-19 patients, its utility is further limited by variability in levels, making exclusion of pulmonary embolism challenging. Miró et al.[28] also found D-dimer less predictive in this group.

The timeline and metabolomic processes underlying pulmonary embolism development in COVID-19 patients still need to be better understood. Additionally, the clinical symptoms of pulmonary embolism are challenging to distinguish, often mimicking acute respiratory distress syndrome (ARDS), a common complication in COVID-19 patients [29]. This symptomatic overlap complicates accurate and timely diagnosis. Developing a comprehensive understanding of the pathophysiological mechanisms of pulmonary embolism in COVID-19 patients, particularly within a temporal or longitudinal framework, is crucial for improving diagnostic accuracy, enabling timely interventions, and ultimately enhancing patient outcomes.

The primary objective of this study is to uncover the metabolic processes that drive pulmonary embolism in COVID-19 patients. It aims to illuminate how COVID-19 amplifies thrombus formation leading to pulmonary embolism and propagation, by investigating the complex interactions between hypercoagulability, endothelial dysfunction, and inflammatory responses reflected in metabolomic profiles.

Coagulation, thrombosis and inflammation drive characteristic shifts in circulating metabolites, ranging from altered amino-acid and lysophospholipid profiles in deep vein thrombosis patients to minor plasma lipids modulating thrombin generation [30,31]. This makes metabolomics suited to capture these pathophysiological signatures. To address these objectives, the metabolomic profile of 66 COVID-19 patients admitted to the Erasmus Intensive Care Unit(ICU) was analysed over 14 days following admission. Targeted LC-MS measurements were conducted on these samples, generating 1209 variables, including individual metabolites (n=1146) and biologically relevant sums and ratios. These features encompassed over 28 lipid subclasses, lipids and amines. We then performed various longitudinal analyses to search for a metabolic pattern that indicates pulmonary embolism.

## 2. METHOD

### 2.1 Study Cohort

The study cohort included 77 adults admitted to the ICU of Erasmus Medical Center, Netherlands, during the initial wave of the COVID-19 pandemic, spanning from February 28, 2020, to April 12, 2020. Among the initial cohort, three patients were excluded due to suspected thrombosis without subsequent confirmation of pulmonary embolism, and one patient was excluded due to the unavailability of Body Mass Index (BMI) data. These exclusions were undertaken to minimise confounding factors and enhance the reliability of the study’s findings. Additionally, seven patients were excluded because their pulmonary embolism diagnoses occurred more than 14 days post-admission to the ICU, attributed to the limited availability of measured samples beyond this period (see Supplementary File **S1** for the study flowchart).

As a result, the final analysis included 66 patients, with 233 blood samples collected within 1–14 days of ICU admission. **Table 1** summarises the demographic and clinical characteristics of the included patients and the collected blood samples. For further details, please refer to the accompanying supplementary information (**Table S1;TableS2**).

**Table 1:**
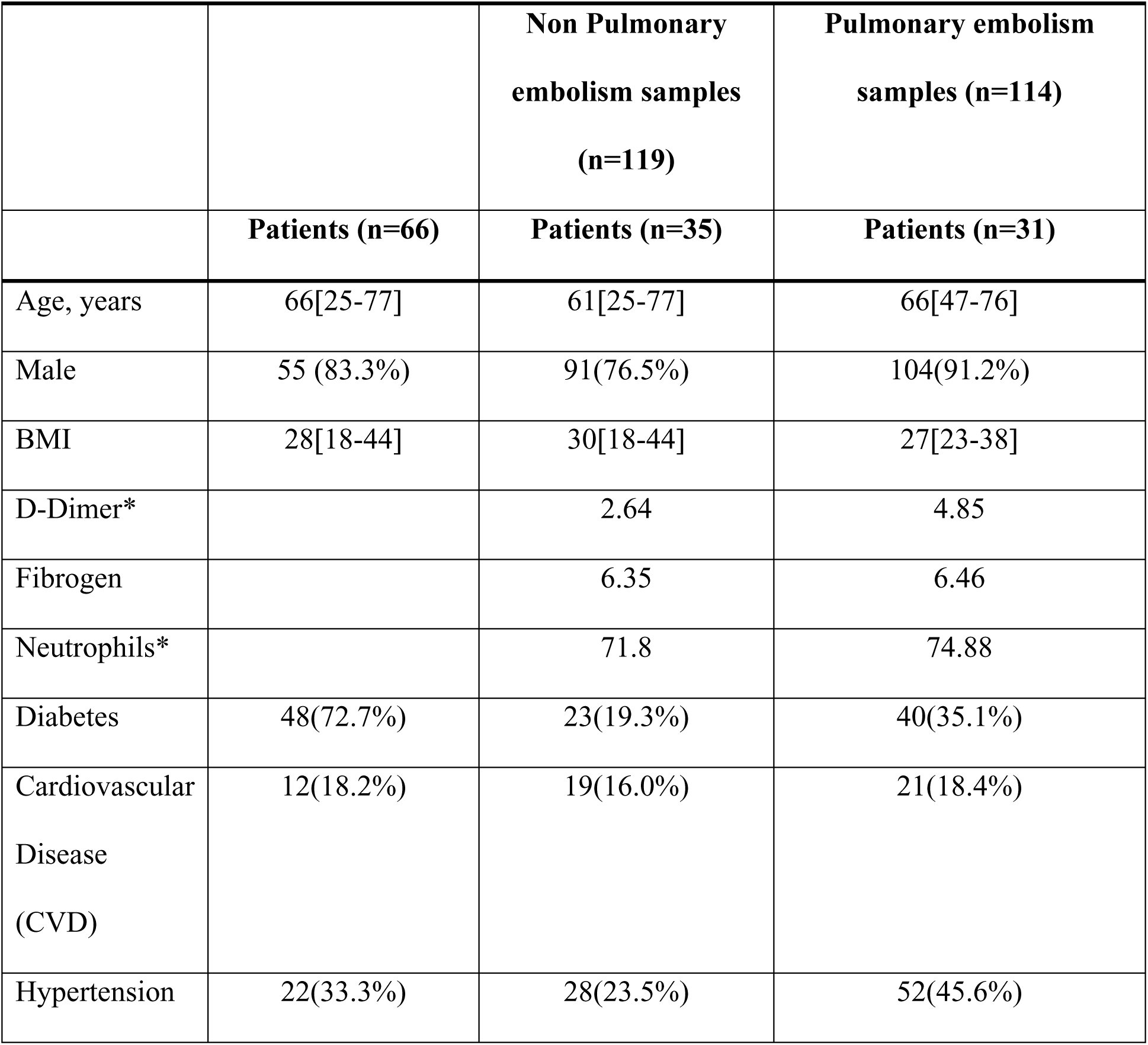
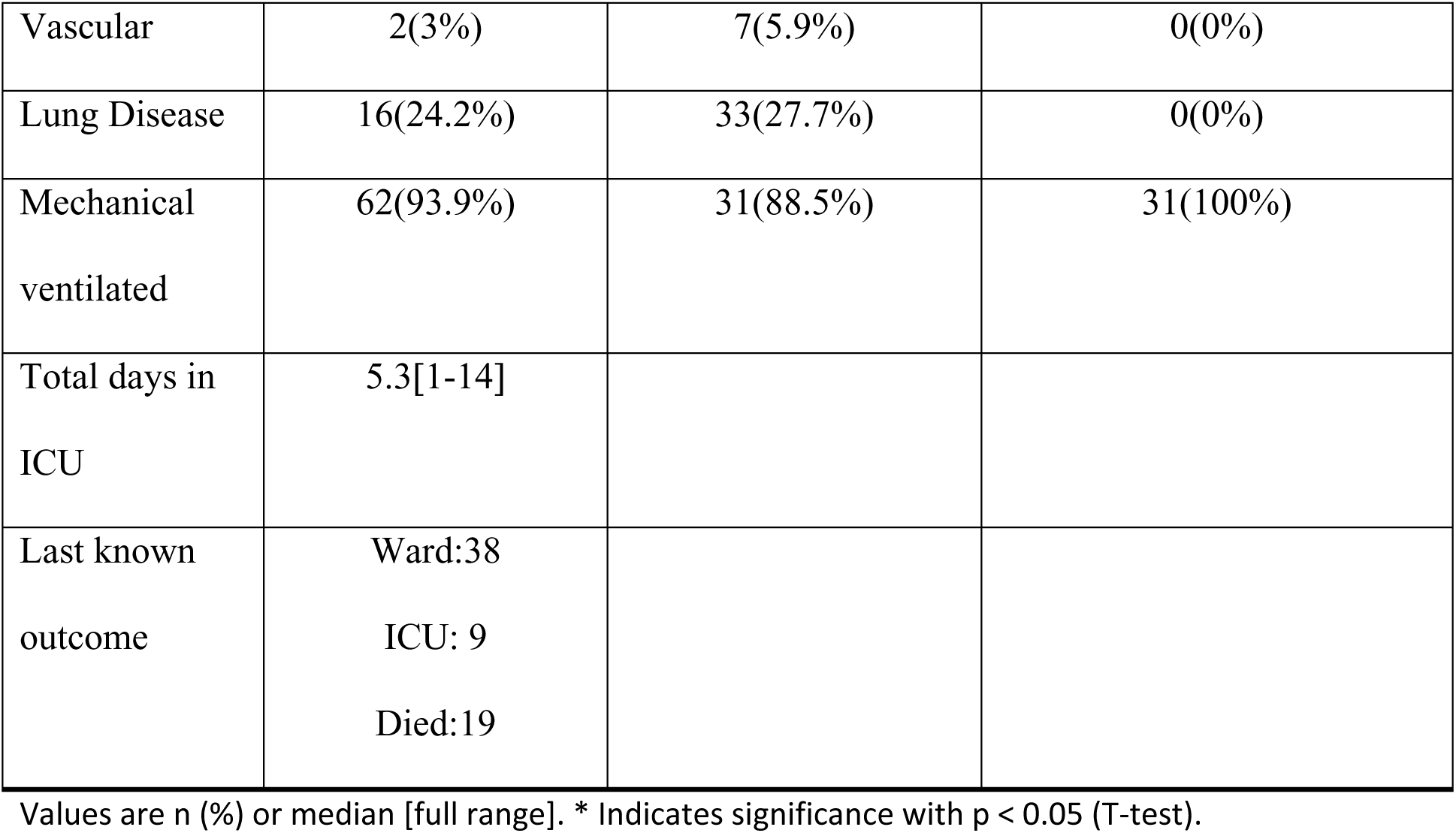
Overview of Selected Demographic and Clinical Features of COVID-19 Samples

|  |  | Non Pulmonary<br>embolism samples<br>(n=119) | Pulmonary embolism<br>samples (n=114) |
| --- | --- | --- | --- |
|  | Patients (n=66) | Patients (n=35) | Patients (n=31) |
| Age, years | 66[25-77] | 61[25-77] | 66[47-76] |
| Male | 55 (83.3%) | 91(76.5%) | 104(91.2%) |
| BMI | 28[18-44] | 30[18-44] | 27[23-38] |
| D-Dimer* |  | 2.64 | 4.85 |
| Fibrogen |  | 6.35 | 6.46 |
| Neutrophils* |  | 71.8 | 74.88 |
| Diabetes | 48(72.7%) | 23(19.3%) | 40(35.1%) |
| Cardiovascular<br>Disease<br>(CVD) | 12(18.2%) | 19(16.0%) | 21(18.4%) |
| Hypertension | 22(33.3%) | 28(23.5%) | 52(45.6%) |
| Vascular | 2(3%) | 7(5.9%) | 0(0%) |
| Lung Disease | 16(24.2%) | 33(27.7%) | 0(0%) |
| Mechanical ventilated | 62(93.9%) | 31(88.5%) | 31(100%) |
| Total days in ICU | 5.3[1-14] |  |  |
| Last known outcome | Ward:38<br>ICU: 9<br>Died:19 |  |  |

Values are n (%) or median [full range]. \* Indicates significance with $p < 0.05$ (T-test).

### 2.2 Samples

EDTA blood samples were obtained at different intervals from 1 to 7 days during the study, as outlined in **Table S2** for each patient. A small portion of the acquired sample was promptly used for flow cytometric immune profiling. Most of the sample was processed to separate the plasma. It was then aliquoted and stored at -20°C until it was ready for serological analysis or transport to the Biomedical Metabolomics Facility Leiden. The plasma samples were maintained at -80°C until further aliquoted and measured using three different LC-MS assays. Refer to the supplementary file **S2** for additional details on the methods on the different LC-MS platforms, including amines, signalling lipids and lipidomics.

### 2.3 Statistical analysis

All data handling and statistical analysis were performed using R v-4.2.1 (see **S3** supplementary file for detailed information). For details on data preprocessing, please refer to supplementary file **S4**. A Principal Component Analysis (PCA) was performed to assess the data variability between the patients who experienced a pulmonary embolism event and those who did not (refer to **S5** supplementary material for PCA details). A correlation analysis was conducted to assess the associations between cytokines and various features. Detailed results of the correlation analysis can be found in supplementary file section **S6** and Table S4.

We compared longitudinal metabolomic profiles between patients who developed pulmonary embolism and those who did not using linear mixed-effects models (*afex* package v1.3-0). Relative metabolite abundance was the dependent variable, pulmonary embolism occurrence was the primary predictor, and models were adjusted for sex, age, BMI, and days since ICU admission. Time (days from ICU admission) was included as a fixed effect, patient ID as a random effect, and a pulmonary embolism × time interaction term tested for group-specific trajectory differences. A patient-level random intercept accounted for repeated sampling (see Supplementary File S7).

The initial analysis assessed overall trajectory differences between pulmonary embolism and non-pulmonary embolism patients. For this study’s aims, three additional group-specific categorical timeframes were defined relative to pulmonary occurrence: (i) 72 hours prior, (ii) 48 hours prior to and including the day of pulmonary embolism, and (iii) 48 hours post and including the day of pulmonary embolism (**Figure 1**). In simple words, we compare metabolite levels in patients with and without pulmonary embolism. First, we look at the average level at the same time point after ICU admission. Then, we check how the levels change over time to see if the groups follow different patterns. The comparisons are made in three ways: between patients with and without pulmonary embolism, between 72 hours before embolism and no embolism, and between 48 hours before plus the day of embolism and no embolism.

**Figure 1:**
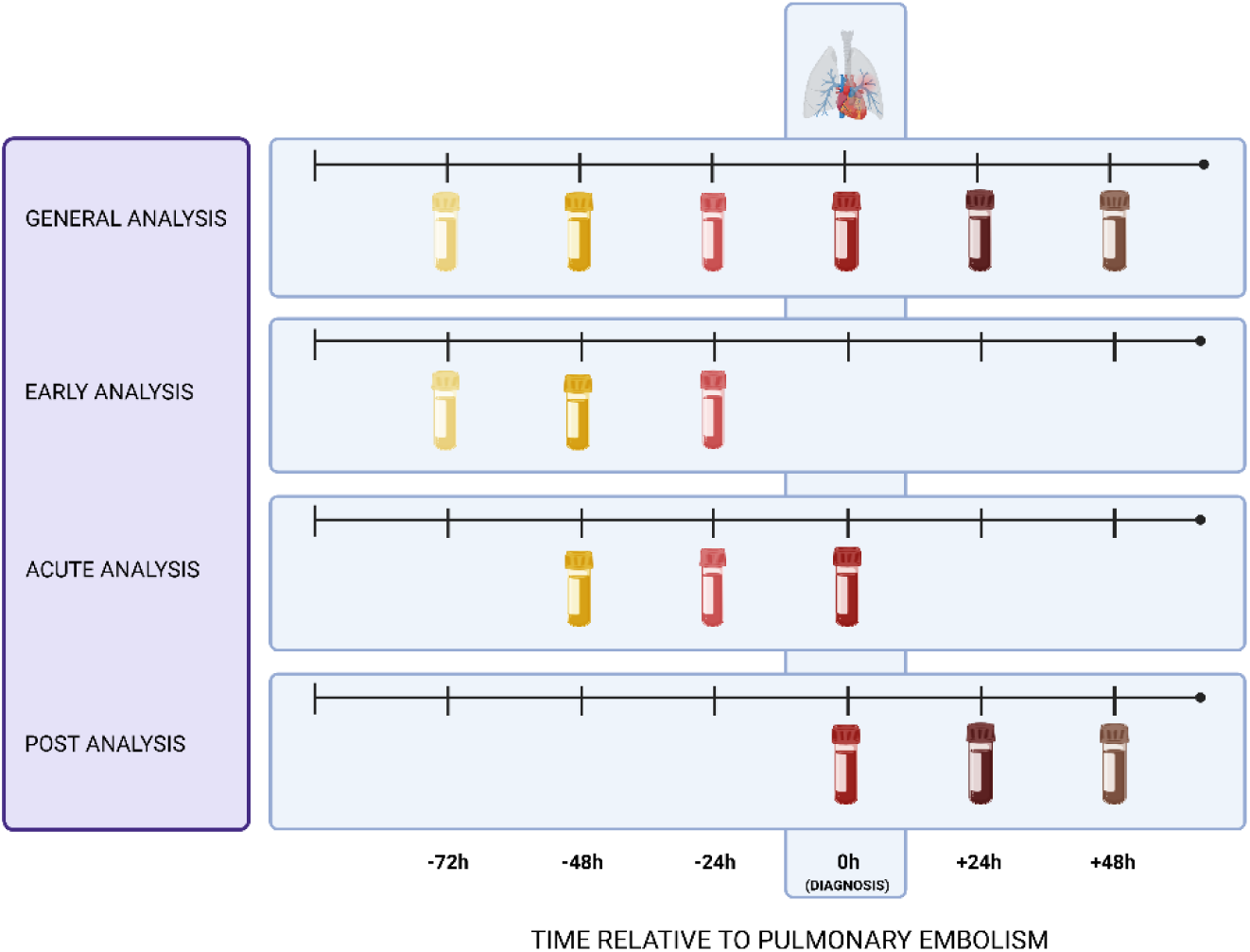
Overview of analyses. Blood sample analyses were conducted in this study, illustrating three distinct timeframes and the overall investigation of pulmonary embolism-associated features, adjusted for age, sex, and BMI. Coloured sections indicate inclusion in the timeframe for comparing pulmonary embolism patients with those without the condition, while grey sections denote exclusion from the temporal comparison analysis. **Created in BioRender.**

The Benjamini-Hochberg method was implemented in the *p*.adjust R function (v-4.2.1) to correct for multiple testing by calculating the false discovery rate adjusted *p*-values (Q-values), where variables with Q < 0.10 were defined as statistically significant.

## 3. RESULTS

### 3.1 Triglycerides and phospholipids increased in patients with pulmonary embolism

The first general comparison was between patients who experienced a pulmonary embolism (n=114 samples) and those who did not (n=119 samples). All samples (n=233) taken from the study patients within 14 days since admission to the Erasmus ICU were included in the general analysis. This analysis identified 115 features with significantly different levels between the two groups at the average time point of 5.5 days post-ICU admission (p-value < 0.05), with three of these features showing significance at Q < 0.1 (**Figure 2a**). In pulmonary embolism, downregulated features included LPE(19:0) and PG(20:3/20:3) with 𝛽_1_ of -0.63 and -0.52 and fold changes of 0.84 and 0.81, respectively, while PG(18:3/20:0) was upregulated (𝛽_1_: 0.64, FC: 1.32) (**Figure 2c**).

**Figure 2:**
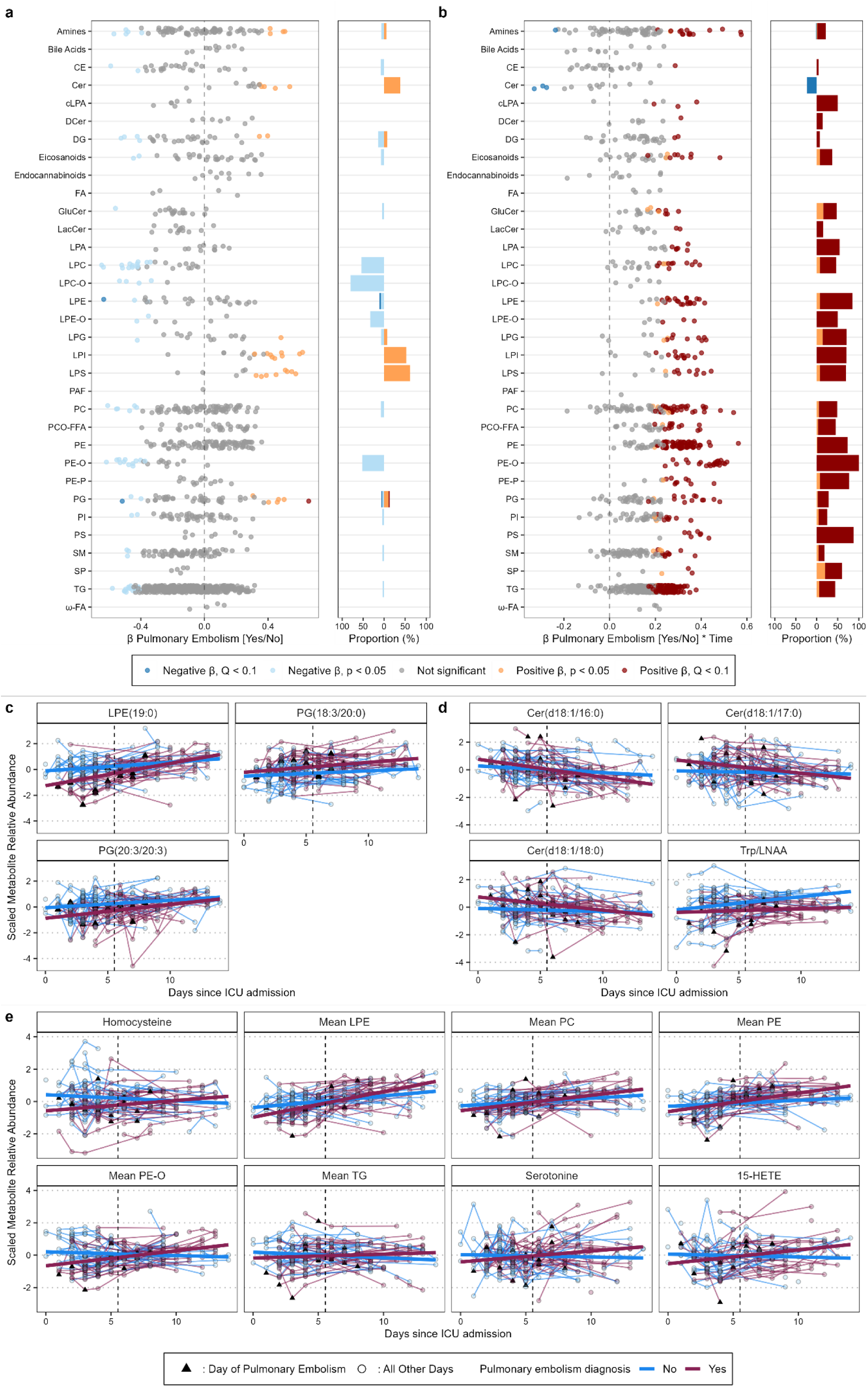
**a)** Associations between features and pulmonary embolism diagnosis. A Linear Mixed Model was utilised, adjusting for days since ICU admission, age, sex, and BMI. Circles indicate the significance of associations, with **grey** representing non-significant associations, while **dark orange** and **dark blue** denote statistically significant positive and negative associations, respectively. Significance was determined using a false discovery rate (FDR) threshold of P < 0.10, adjusted for multiple testing. **b)** The same model as in panel **(a)** illustrates the associations between features and the interaction with days since ICU admission and pulmonary embolism diagnosis, adjusted for the same covariates. **c-e)** Longitudinal trends of significant features identified for COVID-19 patients with (maroon) and without (blue) pulmonary embolism. The vertical dashed line indicates the average day since ICU admission (day 5.3), and the black triangle marks the day of pulmonary embolism for each patient. **(c)** shows primary effect features, with LPE(19:0) exhibiting significant main and interaction effect Q<0.1; (e) shows significant interaction effects

Furthermore, the analysis revealed 464 significant features from 27 classes that showed differential trends over time in patients with pulmonary embolism (Q-value < 0.10, **Figure 2b**). Of the 464 features, four features exhibit a steeper decreasing pattern in pulmonary embolism patients over time. Specifically, three ceramides species (d18:1 with either16:0, 17:0 or 18:0 FA), demonstrated a decline within an estimated range of -0.32 to -0.28, p-values ranging from 0.01 to 0.03, while the Trp/LNAA ratio had an 𝛽_3_ of -0.59, p-value = 0.02, and Q = 0.07) (**Figure 2d**). In contrast, the remaining 460 features showed an increasing trend in pulmonary embolism patients over time, exhibiting a clear class pattern. Remarkably, all 23 ether-linked PE-Os (𝛽_3_ range: 0.25 - 0.51) and 77% of the LPEs were upregulated (𝛽_3_ range: 0.24-0.42) (**Figure 2b**). LPE19:0 was notable for being significantly downregulated in pulmonary embolism patients (𝛽_3_: -0.63, FC: 0.84) but showing a positive trend over time (𝛽_3_: 0.36, **Figure 2c**).

Of the analysed amines (n=74), 21.6% had a steeper increase in pulmonary patients over time. These included alanine, cysteine, homocitrulline, methionine, methylcysteine, serine, serotonin, tryptophan, serotonin/Trp and carnosine (𝛽_3_ range: 0.23 - 0.57; **Figure 2e**). Within the analysed signalling lipids, nearly one-third of all analysed eicosanoids were found significant. These include 11,12-DiHETrE, 9-HOTrE, 13-HODE, 14,15-DiHETE, 15-HETrE,15-HETE and 9,10-DiHOME (𝛽_3_ range: 0.088 - 0.15; p < 0.03, Q < 0.10) (**Figure 2e)**.

### 3.2 Subtle metabolic dysregulation 72 hours prior to pulmonary embolism event

To investigate the pathophysiological processes and metabolic alterations contributing to the onset of a pulmonary embolism event, and to identify potential diagnostic biomarkers predicting a pulmonary embolism event occurring within the next 72 hours, we analysed blood samples from the 72 hours prior to the event (n= 34 samples). This analysis excluded the actual day of pulmonary embolism to gain insights into potential factors preceding the event without interfering with any metabolic changes occurring during the event or as a result of medicinal intervention. Examining the main impact of the 72 hours preceding the occurrence of pulmonary embolism (averaged over the ICU hospitalisation), revealed differential abundances in 101 features across 23 classes (**Figure 3a**). Four features remained statistically significant after applying multiple comparisons correction. Of these four, PC(19:0/20:4) (𝛽_1_ = -0.93), LPC(22:4) (𝛽_1_ = -0.81), and LPE(19:0) (𝛽_1_ = -0.76) were downregulated, while Cer(d18:1/26:1) was upregulated in pulmonary embolism patients (estimate = 0.68; **Figure 3c)**.

**Figure 3:**
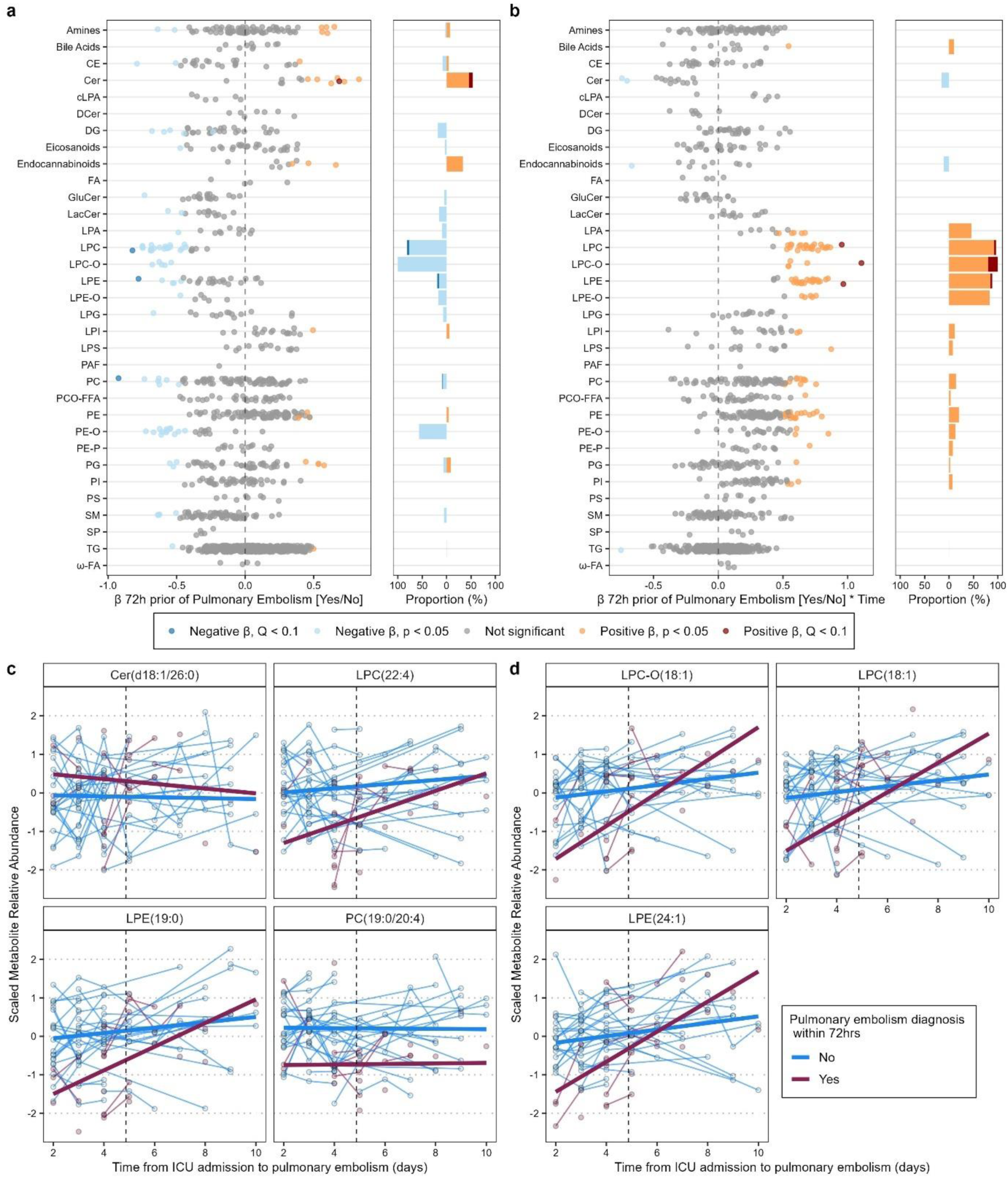
**a)** Associations between features and 72 prior to pulmonary embolism. A linear mixed model was performed by adjusting the number of days since ICU admission, age, sex, and BMI. The 72h prior to pulmonary embolism coefficient of each species is shown on the x-axis. Circles: Gray denotes no significant associations; dark orange and dark blue represent positive and negative significant associations adjusted for multiple testing (P < 10% FDR) **b)** Same model as in panel **(a)**, showing associations between features and the 72 hours prior to pulmonary embolism, along with its interaction with days since ICU admission, adjusted for the same covariates; Longitudinal trends of significant main effects of lysophosphatidyls and ceramide (**c**), and significant interaction effects of lysophosphatidyls (**d**) in the early analysis in COVID-19 patients with (maroon) and without (blue) pulmonary embolism.

Next, we compared longitudinal profiles of blood from patients diagnosed with a pulmonary embolism within 72 hours and those who were not. This analysis revealed 110 features from 18 classes with differential abundances over time (p-value < 0.05; **Figure 3b**). Of these, LPC-O(18:1), LPC(18:1) and LPE(24:1) were significantly upregulated 72hrs prior the pulmonary embolism event (𝛽_3_ range= 0.28-0.32; **Figure 3d**).

### 3.3 Increased PEs within 48 hours and on the day of pulmonary embolism diagnosis

To evaluate the metabolic changes occurring between the onset of pulmonary embolism and its diagnosis, an additional analysis was conducted, encompassing the 48-hour period prior to the diagnosis and the day of the pulmonary embolism diagnosis (n=35 samples). The goal was to identify key biochemical or physiological alterations that could indicate the imminent development of pulmonary embolism during this period. Additionally, this analysis aimed to explore the role of other features changed on the day of pulmonary embolism, in contrast to the previous analysis where this day was excluded.

First, we evaluated the main effect of pulmonary embolism 48 hours before the event and on its day. Here, we identified 106 features with differential abundances, none of which remained significant after correction for multiple comparisons (**Figure 4a**). However, LPE19:0 showed the same downregulation trend as in the previous two analyses.

**Figure 4:**
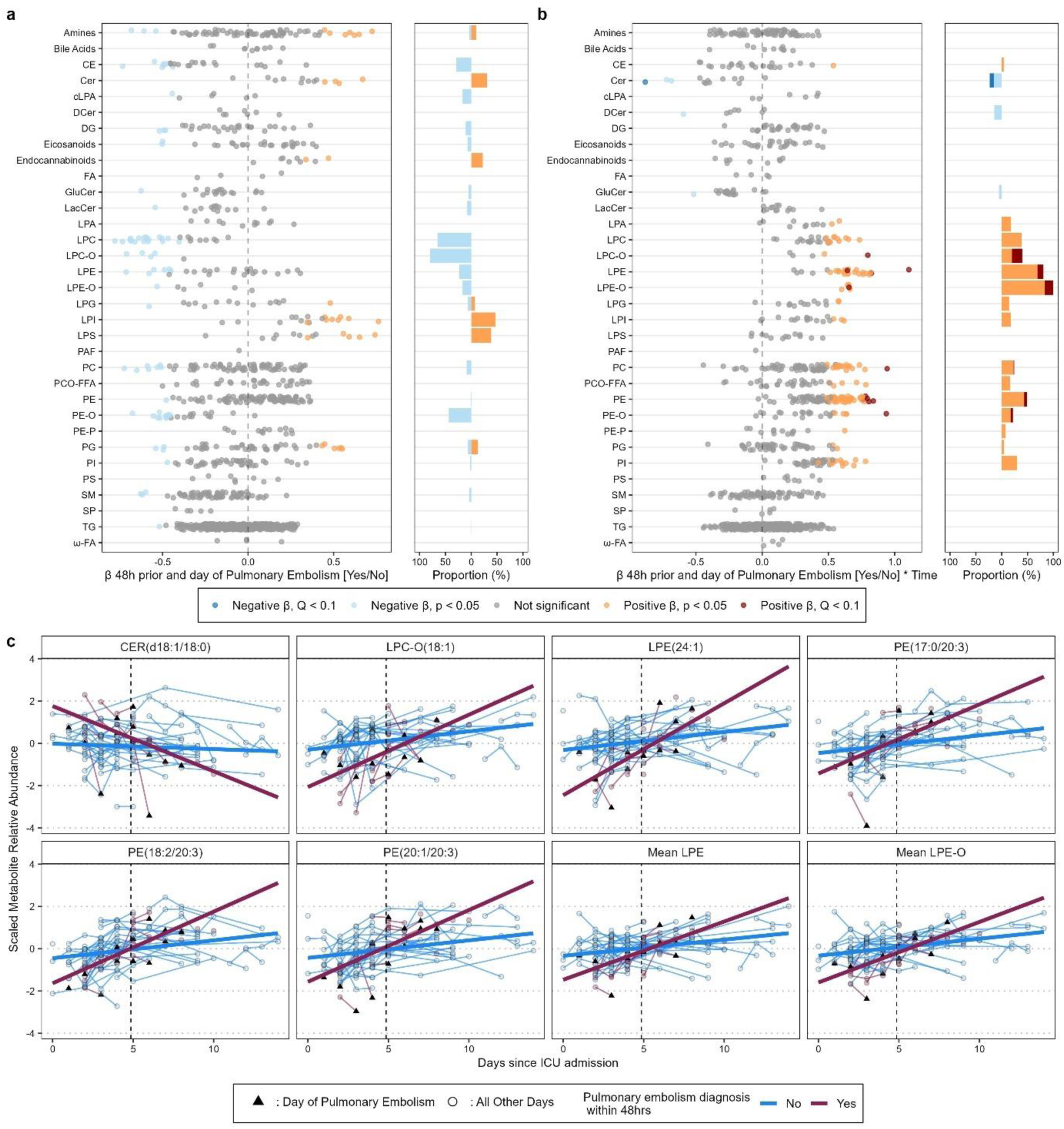
**a)** Associations between features and 48 prior to pulmonary embolism. A Linear Mixed Model was performed by adjusting for days since ICU admission, age, sex, and BMI. The 48h prior to pulmonary embolism coefficient of each species is shown on the x-axis. Circles: Gray denotes no significant associations; dark orange and dark blue represent positive and negative significant associations adjusted for multiple testing (P < 10% FDR) **b)** Same model as in panel **(a)**, showing associations between features and the 48 hours prior to pulmonary embolism, along with its interaction with days since ICU admission, adjusted for the same covariates. **c)** Longitudinal trends of significant interaction effects of lysophosphatidyls, ceramide, and phosphatidylethanolamines identified in the acute analysis among COVID-19 patients with (maroon) and without (blue) pulmonary embolism. The vertical dashed line represents the average day since ICU admission (day 4.87), and the black triangle indicates the day of the pulmonary embolism per patient.

Next, we examined the interaction effect of pulmonary embolism during the 48 hours prior to the event, the day of the event, and the duration since ICU admission. Here, a total of 13 features from seven distinct classes were significantly changed, after multiple testing correction (Q < 0.10) **(Figure 4b)**. PEs were most represented (38.5%; 𝛽_3_: 0.76-0.83; p< 2×10 ^−16^, Q <0.09), and remarkably, 3 of the 4 upregulated PEs contained a side chain with 20:3 fatty acid side chain ((17:0/20:3), (18:2/20:3), (20.1/20.3) and a PE-O(18.0/20.3)). Furthermore, two LPEs (LPE(14:0) and LPE(24:1)) were upregulated over time (𝛽_3_: 0.82 - 1.11; p < 2×10 ^−16^; Q = 0.08) alongside the mean aggregated z-score of the LPE-Os (𝛽_3_: 0.66; p < 2×10 ^−16^, Q < 0.10) as well as the mean aggregated z-score of the LPEs (𝛽_3_: 0.63; p < 0.001, Q < 0.10). Only Cer(d18:1/18:0) showed a negative trend over time (𝛽_3_ =-0.88; p-value = 0.001, Q = 0.10) (**Figure 4c**). Both LPC-O(18:1) and LPE(24:1), which were significantly upregulated 72 hours prior to the pulmonary event, were also significant in this analysis after FDR correction (𝛽_3_ = 0.81, p-value = 0.001, Q = 0.09).

### 3.4 Strong triglyceride and phosphoethanolamine signature during and post pulmonary embolism

To understand the metabolic changes in the period covering the day of the pulmonary embolism diagnosis, we analysed the interval of the day of diagnosis and the following 48 hours.

First, we compared the day of pulmonary embolism and the subsequent 48 hours to periods while adjusting for the average number of ICU days (n=33 samples). The initial results showed 186 features with differing levels between patients with and without embolism (**Figure 5a)**. Of these 55 features from 12 classes remained significant after multiple testing correction with Q < 0.1, including 11 elevated lysophosphatidylinositols (LPIs), representing 65% of all measured LPIs, with estimates ranging from 0.74 to 1.38 and fold changes ranging from 1.44 to 2.64 (**Figure 5c**). The LPIs predominantly had polyunsaturated FAs (80.0%), with long-chain FAs (≥C20) representing the largest fraction (40.0%) and monounsaturated FAs comprising only 20.0%.

**Figure 5:**
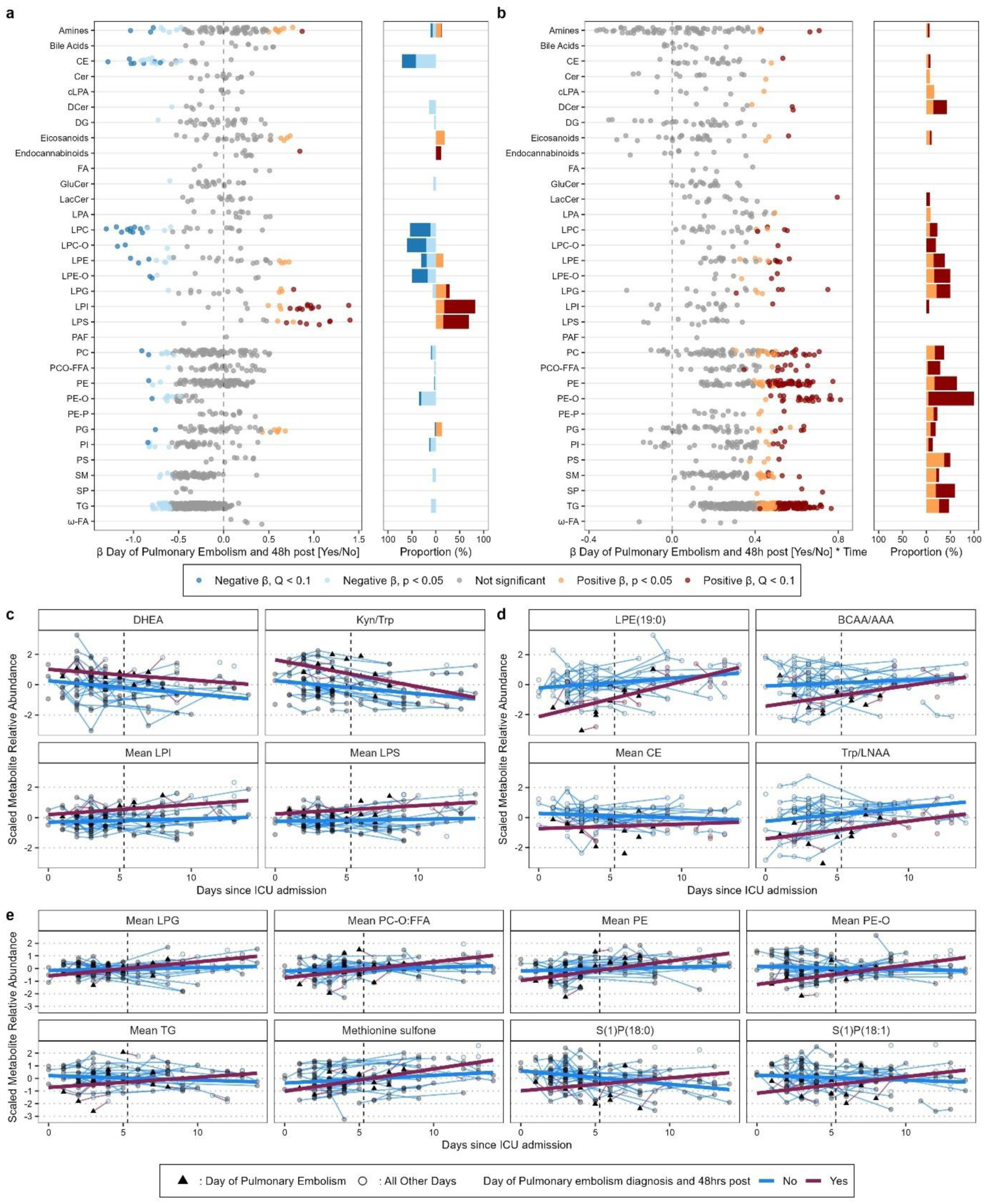
**a)** Associations between features and the 48-hour period following pulmonary embolism. A Linear Mixed Model was utilised, adjusting for days since ICU admission, age, sex, and BMI. The coefficients for each feature 48 hours post-pulmonary embolism are plotted on the x-axis. Circles indicate the significance of associations, with grey representing non-significant associations, while dark orange and dark blue denote statistically significant positive and negative associations, respectively. Significance was determined using a false discovery rate (FDR) threshold of P < 0.10, adjusted for multiple testing. **b)** The same model as in panel **(a)** illustrates the associations between features during the 48-hour period following pulmonary embolism, including their interaction with days since ICU admission, adjusted for the same covariates. **c)** Longitudinal trends of significantly upregulated features identified in the post-analysis for COVID-19 patients with (maroon) and without (blue) pulmonary embolism. The vertical dashed line indicates the average day since ICU admission (day 5.3). **d)** Longitudinal trends of significantly downregulated features identified in the post-analysis for COVID-19 patients with (maroon) and without (blue) pulmonary embolism. The vertical dashed line indicates the average day since ICU admission (day 5.3). **e)** Longitudinal trends of significant interaction effects of lysophosphatidyls, triglycerides, methionine sulfone, sphingosine, and phosphatidylethanolamines identified in the post-analysis among COVID-19 patients with (maroon) and without (blue) pulmonary embolism. The vertical dashed line is the average day since ICU admission (day 5.3), and the black triangle marks the day of pulmonary embolism for each patient.

This was followed by decreased LPCs(n=11), with 𝛽_3_ranging from -1.31 to -0.86 and FCs from 0.64 to 0.83, representing 42% of the LPCs. The significant LPCs are predominantly with saturated FAs (81.8%), with long-chain FAs (≥C19) representing the largest fraction (45.5%) and monounsaturated FAs (18.2%) restricted to the long-chain category (LPC 20:1, LPC 24:1). Additionally, 12% belonged to lysophosphatidylserines (LPSs, n=7), which were also elevated with estimates ranging from 0.71 to 1.39 and fold changes ranging from 1.57 to 3 representing 53.8% of the LPSs. The significant LPSs are all polyunsaturated LPSs, with long-chain LPSs (≥C19) representing the largest fraction (83.4%) (**Figure 5c**).

When looking into the interaction effect between the day of the pulmonary embolism diagnosis and the 48 hours immediately thereafter and the days since admission to ICU, 223 features (Q < 0.10) covering 23 metabolite classes were significantly different (**Figure 5b**). Of these, 41% were upregulated TGs (n=91, 𝛽_3_: 0.41-0.78), representing a fifth of all measured TGs.

Remarkably, 95.7% of all ether-linked PE-Os (𝛽_3_ range: 0.47 - 0.79) and 46.8% of the PEs were upregulated (𝛽_3_ range: 0.45-0.76) (**Figure 5b**).

Additionally, two sphingosine-phosphate species (40%) were upregulated, with S1P18:0 displaying a β₃ value of 0.739 and S1P(18:1) a β₃ value of 0.61(**Figure 5e**).

Three amino acids metabolites were found to be significantly elevated, namely methionine sulfone (𝛽_3_ = 0.42, p-value = 0.015, Q = 0.09), taurine(𝛽_3_ = 0.70, p-value = 0.001, Q = 0.04) and carosine(𝛽_3_ = 0.65, p-value = 0.002, Q = 0.06).

## 4. DISCUSSION

### 4.1 Summary and overall altered chemical classes

The results detailed in our study are discussed (per analysis), within their metabolic and biochemical context, incorporating findings from relevant studies and providing hypotheses for the observations.

The analysis revealed the greatest overlap of features between the general analysis and the period spanning the day of pulmonary embolism and up to 48 hours post-event. This overlap encompassed 163 metabolite features, with the majority classified as TGs (n=46), followed by PEs (n=42), PE-Os (n=22), and PCs (n=14) (**Figure 6**). A secondary overlap was observed among 10 features shared between the general analysis, the day of pulmonary embolism, and 48 hours preceding the event. Among these, six were were PEs, including three LysoPEs. Notably, both LPC-O(18:1) and LPE(24:1) features were identified as overlapping in both early-stage and acute-phase analyses.

**Figure 6:**
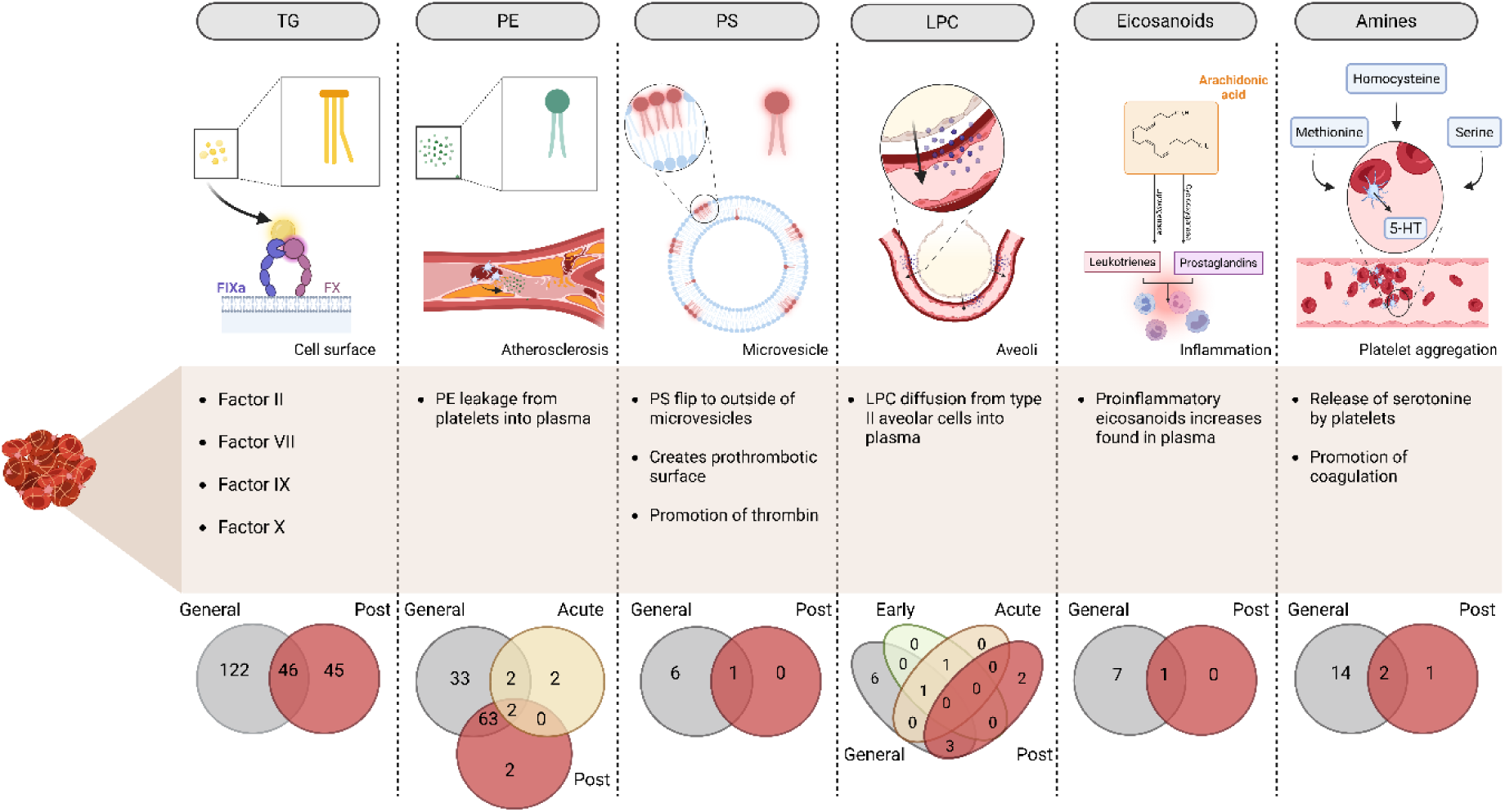
Summary of Key Analytical Findings. The diagram highlights the most significantly altered metabolite classes found in the analysis’s, categorised by their partial functional roles, with hypotheses regarding their involvement in thrombosis /pulmonary embolism. Below each class a Venn diagram presents data from different time points, showcasing temporal changes in metabolic activity and overlaps between these classes over time. **Created in BioRender.**

### 4.2 Metabolic Changes generally Associated with Pulmonary Embolism

The marked increase in TGs and phospholipids may reflect the body’s response to pulmonary embolism’s inflammatory and pro-thrombotic state [32–35]. Our findings indicate that TGs significantly increase in patients who develop pulmonary embolism. These were among the 464 feature that significantly changed (39.2%), suggesting a potential lipid-related mechanism that contributes to thromboembolism in COVID-19. These results are consistent with epidemiological studies that demonstrate an association between hypertriglyceridemia and elevated levels of vitamin K-dependent procoagulant factors, which are known to bind triglyceride-rich lipoproteins and enhance procoagulant reactions (**Figure 6**)[36–41]. Such a relationship may underpin a prothrombotic state in individuals with elevated TGs, thereby increasing the risk of pulmonary embolism in this subpopulation.

The observed elevation in TGs may be pivotal in promoting thrombosis through multiple pathways. In addition, hypertriglyceridemia is also associated with pathological platelet hyperactivation [42,43]. This hyperactivation can lead to the release of platelet microvesicles and granules, further amplifying the pro-thrombotic state. Consequently, this may increase the risk of thrombosis and trigger a cytokine storm, both at local and systemic levels [44]. The formation of platelet-leukocyte conjugates, particularly with neutrophils, and the subsequent processes of platelet apoptosis or aggregation further exacerbate thrombotic and inflammatory responses, which are key pathological features in pulmonary embolism [17,18,44,45].

Additionally, impaired fibrinolysis and dysfunction of endothelium associated with increased TGs in blood, can exacerbate thrombosis leading to life-threatening thromboembolic events [46]. The lungs are particularly vulnerable to such thrombotic processes, which could result in the development of a pulmonary embolism. Autopsy studies in patients with COVID-19 have highlighted the presence of thrombosis in both arterial and venous vessels, especially in the form of platelet-rich thrombotic microangiopathy within small vessels [18,44]. This pathology has been associated with ARDS and diffusely oedematous lungs, suggesting that hyperactivated platelets not only contribute to thrombosis but also exacerbate pulmonary inflammation [44]. However, it remains challenging to determine the precise extent and contribution of elevated circulating lipids due to tissue and cellular damage. Disease progression may lead to the release of catabolites and metabolic byproducts into the bloodstream, further complicating this assessment.

Phosphatidylethanolamines (PEs) were mentioned as potential accelerators of blood coagulation, either independently or combined with other phospholipids [47]. However, other studies have challenged these findings, which may be attributed to variations in experimental conditions, such as differences in the phospholipid particles’ colloidal state, pH levels, ionic strength, and purity[48]. Despite these conflicting results, there is general agreement that platelet function in coagulation is closely linked to their phospholipid content. Notably, an increase in PE levels has been observed in the plasma of patients with chronic arterial occlusive disease and ischemic heart disease, suggesting its potential relevance in the progression of these conditions that could also be linked to pulmonary embolism. The study of Bruhn et al.[49] showed a significant decrease in PE levels in the platelets of patients with chronic arterial occlusions compared to healthy individuals. The exact reason for this decrease is unclear, but it is hypothesised that PEs may leak from platelets into the plasma in patients with atherosclerosis (**Figure 6**). This observation is particularly intriguing because such leakage appears to be specific to PEs, as other lipids do not exhibit similar differences between healthy and diseased states [49]. Notably, PEs (14.5%) demonstrated significant upregulation, particularly during the acute phase of pulmonary embolism. The presence of a 20:3 fatty acid side chain in 7 of 68 upregulated PEs suggests a specific lipid remodelling process, which may be critical in the pathogenesis of pulmonary embolism, as discussed above. Furthermore, LPEs, especially LPE(19:0) and LPE(24:1), were significantly upregulated over time and found in the general and early phase of pulmonary embolism diagnosis. These two represent interesting lipid classes of odd-chain fatty acids, and very-long chain fatty acids (VLCFAs). This may suggest a link to peroxisomal dysfunction (decrease in peroxisomal and numbers) that exacerbates mitochrondrial oxidative stress, increases lipid peroxidation, and affects the metabolism of lipids such as ether-linked phospholipids, bile acids, VLCFAs (cannot be metabolised in the mitochondria), and DHA (FA 22:6). The resulting lipid dysregulation and downstream effects may compromise the innate immune response and inflammation signalling.

Notably, levels of LPCs and LPEs were significantly elevated in the blood of COVID-19 patients at high risk of thrombosis compared to those at lower risk (D-dimer ≤ 900 U/mL) [50].

Phosphatidylserine (PSs; 4.3%) also demonstrated a substantial upregulation, with 87.5% of the PSs being significant. PSs is one of the most abundant phospholipids in plasma membranes. Under normal conditions, it is primarily found on the inner side of the membrane [51,52]. This positioning is regulated by an ATP-dependent enzyme called *flippase*, which swiftly moves any PSs that migrate to the outer membrane back to the inside of the cell [51,52]. However, when cells are activated or injured, as seen in critical illnesses like COVID-19,[52] . PSs can become exposed on the outer membrane surface, allowing coagulation proteins to assemble on what was once a non-thrombogenic surface. Microvesicles that are released into the plasma after exocytosis also portray this behaviour. (**Figure 6**) [51–55]. This interaction within the coagulation cascade ultimately leads to thrombin formation.

The phosphatidylcholines (PCs; 7.3%) showed an upward trend in patients with pulmonary embolism. This aligns with previous research suggesting lipid metabolism is often disrupted in critical illnesses and inflammatory conditions such as COVID-19 [56].

The upregulation of eicosanoids such as 14,15-DiHETE, 9,10-DiHOME, 11,12-DiHETrE, 13-HODE and 15-HETE in COVID-19 patients who experienced pulmonary embolism suggests a complex interplay of inflammatory and vascular responses (**Figure 6**) [57,58]. Eicosanoids, derived from polyunsaturated fatty acids through various enzymatic pathways, play crucial roles in inflammation, immunity, and cardiovascular health. For instance, 11,12-DiHETrE, derived from arachidonic acid by CYP epoxygenase followed by sHE hydrolysis is involved in anti-inflammatory responses and vasodilation [57–59].

The upregulation of pro-inflammatory eicosanoids like 9,10-DiHOME, 13-HODE, and 15-HETE indicates an enhanced inflammatory state associated with COVID-19 and pulmonary embolism. The two linoleic acid-derived products 9,10-DiHOME, (produced via CYP and sEH) and 13-HODE (via 15-LOX enzyme), both act as inflammatory mediators contributing to endothelial damage and hypercoagulability in COVID-19 patients [60–62].

Furthermore, 15-HETE produced from arachidonic acid through 15-LOX is particularly notable for its suspected contributions to endothelial dysfunction,[63,64] platelet activation,[65,66] and the amplification of the coagulation cascade,[67] all of which are critical in the pathogenesis of pulmonary embolism[68,69].

14,15-DiHETE (from EPA) and 11,12-DiHETrE (from AA) play a role in the regulation of vascular tone and inflammation [70]. Notably, 11,12-DiHETrE exhibited a negative correlation with IL-6 (ρ = -0.40, Q = 0.03) and neutrophil count (ρ = -0.44, Q = 0.05) in patients with pulmonary embolism. These findings support the proposed role of 11,12-DiHETrE as an anti-inflammatory biomarker.

COVID-19-associated coagulopathy (CAC) exemplifies a mechanism which involves complex, dysregulated interactions between various clotting and immune systems, including the innate and adaptive immune systems [17,56,69,71]. This integration of innate immune responses with coagulation, especially within the microvasculature where endothelial cell dysfunction promotes clot formation and inflammation, is termed *immunothrombosis.*[12,13,17,18]. The primary difference between CAC and conventional clotting disorders lies in their mechanisms. Conventional clotting issues rely heavily on thrombin production, whereas CAC results from inflammation, endothelial dysfunction, platelet activation, and immune response abnormalities. The SARS-CoV-2 spike protein further influences the clotting process, contributing to CAC’s unique characteristics [18,69,71].

The significant upregulation of 16 amines, including homocysteine, serotonin, methionine, serine and the serotonin/Trp ratio, suggests their involvement in stimulating platelet aggregation [72,73]. This is in line with the previously reported hypercoagulable state in these patients, which could contribute to the development of pulmonary embolism [17,18,69,71,74]. Elevated levels of homocysteine and serotonin, are known to promote platelet aggregation and endothelial dysfunction, aligned with the increased risk of thrombotic events observed in COVID-19 (**Figure 6**) [73,75]. Similarly, the upregulation of methionine and serine, both involved in the methylation processes and metabolic pathways negatively influencing platelet function, further supports the prothrombotic milieu [73].

The combined upregulation of these features and amines suggests a multifaceted pathophysiological mechanism involving both inflammatory and coagulation pathways, providing an understanding of the heightened thrombotic risk in COVID-19 patients with pulmonary embolism.

### 4.3 Temporal Dynamics of Metabolic Alterations

Analysing the metabolic changes at different time points relative to the pulmonary embolism diagnosis provided insights into the temporal dynamics of these alterations.

LPC-O(18:1), LPC(18:1), and LPE(24:1) significantly increase within 72 hours before pulmonary embolism, indicating that these metabolites may serve as early predictive biomarkers for thromboembolic events in COVID-19 patients. These changes may reflect early perturbations in lung surfactant metabolism, which is dominated by phospholipids, approximately 80% PCs and a smaller proportion of LPCs. Alterations in lung surfactant composition, including variations in total phospholipid content and specific surfactant phospholipid classes, have been observed in pulmonary embolism [76]. These modifications likely result from cellular stress induced by hypoxia, reactive oxygen species, and plasma protein leakage into the alveoli [77].

Damage to alveolar type II cells increases membrane permeability, potentially leading to the leakage of surfactant components into both the bloodstream and alveolar spaces (**Figure 6**) [77]. Previous studies have confirmed the presence of surfactant-specific proteins, namely SP-A and SP-D, in the blood, reinforcing the notion that surfactant components can serve as biomarkers for lung injury [78]. Consequently, elevated plasma levels of PCs—and to a lesser extent, LPCs—may serve as early indicators of lung injury associated with pulmonary embolism [76,79,80].

Dipalmitoylphosphatidylcholine (DPPC) is the primary surface-active phospholipid in lung surfactant [81]. However, in this study, DPPC levels did not exhibit significant changes. In contrast, PC(16:0/18:1), another phosphatidylcholine species specific to lung surfactant[81], showed a statistically significant increase. While these findings suggest a potential role for PCs and LPCs as biomarkers of pulmonary embolism-related lung injury, they should be interpreted with caution. The observed changes may be influenced by multiple physiological and pathological factors, and their specificity to pulmonary embolism remains uncertain. Further rigorous investigation, including larger studies and mechanistic analyses, is essential before any definitive conclusions can be drawn regarding their clinical utility.

The increase in ceramide(26:1) 72hr prior to embolism may suggest involvement of peroxisomal dysfunction. Also, long-chain ceramides can permeabilize and damage mitochondria, leading to a decrease in electron transport chain activity and compromise energy production [82].

Within the 48 hours following and including the day of pulmonary embolism diagnosis, we observed a significant upregulation of PEs, particularly those containing the 20:3 fatty acid side chain. This period also showed increased levels of LPEs and a notable upregulation in the mean aggregated z-score of LPE-Os. These findings underscore the potential for these lipid features to serve as immediate biomarkers for the onset of pulmonary embolism [56].

### 4.4 Metabolic Signature During and Post Pulmonary Embolism Diagnosis

Elevated triglyceride levels have been increasingly recognised in the literature as a potential risk factor for the development of pulmonary embolism, primarily through their association with deep vein thrombosis [35,83–85]. Some studies have challenged this claim, suggesting that factors such as obesity and smoking were more strongly associated [86,87]. Several studies suggest that hypertriglyceridemia contributes to the hypercoagulable state by altering lipid metabolism, which increases blood viscosity and enhances the risk of thrombus formation in the deep veins, as discussed in section **4.1**. This study found that triglyceride levels were significantly elevated; TGs accounted for 40.8% of the features. This elevation in TGs during and post-pulmonary embolism suggests that while high TG levels may predispose individuals to deep vein thrombosis and subsequent pulmonary embolism, they are notably more pronounced during or after the embolic event, highlighting their role not only in the risk of thrombus formation but also as a marker of embolic severity.

This strong TG signature supports the hypothesis that TG metabolism is significantly altered during acute pulmonary embolism events, potentially due to increased lipolysis and lipid mobilisation as part of the body’s acute phase response [32].

Phospholipase A2 (PLA2) activity is central to lipid metabolism, particularly in response to cellular stress and tissue damage. PLA2 hydrolyses membrane phospholipids, releasing bioactive lipids such as arachidonic acid and LPA, a pro-thrombotic mediator that increases the risk of thrombosis [56]. In line with this, the study revealed a significant upregulation of PEs, PE-Os, and PCs during and immediately after a pulmonary embolism event. The substantial representation of these lipid classes further supports the notion of broad lipid metabolism dysregulation. This finding is consistent with studies on COVID-19, where increased PEs and PCs were linked to PLA2 activity, reinforcing the connection between lipid alterations, inflammation, and thrombotic risk [56].

The kyurenine-to-tryptophan (Kyn/Trp) ratio has emerged as an important biomarker of both acute and chronic inflammation. Elevated levels of kyrurenine as well as this ratio have been widely reported in various diseases, including COVID-19, and other inflammatory conditions [88–93]. Since kyrurenine is formed via inflammation-triggered enzyme IDO and induces further inflammation this ratio serves as an indicator of immune activation, reflecting the degree of inflammation and its potential impact on disease progression. In line with this definition, we found Kyn/Trp was also positively correlated with IL-6(𝜌=0.62, Q=0.01) and neutrophils (𝜌=0.49,Q < 2×10 ^−16^). Its significance in COVID-19 underscores the broader role of tryptophan metabolism in immune regulation and suggests potential avenues for therapeutic intervention.

Methionine sulfone, a strong marker of oxidative stress, arises from the post-translational modification of the methionine residue in proteins [94]. Its presence has been noted in COVID-19 and other lung diseases, highlighting its relevance in the disease pathophysiology that involves both oxidative stress and inflammation [88].

Sphingoid base 1-phosphates (S1Ps) are key immune modulators involved in immune cell trafficking and endothelial function, regulated by specific cell surface receptors [95]. The S1P 18:1 metabolite plays a central role in lymphocyte movement between lymphoid organs and the bloodstream, and is also linked to inflammation resolution in conditions like acute lung injury. In the context of COVID-19 and pulmonary embolism, lower levels of certain S1P metabolites are associated with severe disease, suggesting their potential as prognostic markers for disease outcomes [95–97]. In our study, we find that S1P 18:0 and S1P 18:1 have lower baseline levels in pulmonary patients, but increase in a steeper trend over time, perhaps due to treatment.

A limitation of our study was the absence of a true control group, as all patients included were COVID-19 positive and admitted to the ICU. The findings may not be broadly generalisable beyond this specific patient population. However, this focus strengthens the study by providing in-depth insights into the critical care needs of severe COVID-19 cases, offering valuable clinical guidance for managing similar future outbreaks or respiratory infections requiring intensive care. Another important limitation is the uncertainty regarding the exact timing of pulmonary embolism onset. Our data only include the time of pulmonary embolism diagnosis, not the precise moment when the event occurred. However, given the acute nature of pulmonary embolism and its rapid clinical progression, we expect that the time between embolism occurrence and diagnosis was relatively short. Nonetheless, this uncertainty adds complexity to the interpretation of biomarker changes and their potential as early indicators of lung injury.

## 5 CONCLUSION AND IMPLICATIONS FOR CLINICAL PRACTICE

In this study, we studied metabolic changes in COVID-19 patients with and without pulmonary embolism and found a clear connection between higher lipid levels and the development of embolic events. When we tracked patients over time, we saw subtle shifts in metabolism before pulmonary embolism occurred, suggesting that these subtle changes could signal an increased risk. These early signals need to be confirmed in larger, forward-looking studies, but they point to lipid metabolism as an important area for both diagnosis and treatment. In particular, certain lipid markers, such as LPC-O(18:1) and LPE(24:1), showed strong potential as early warning signs for pulmonary embolism. This highlights the value of lipid profiling not only for assessing risk but also for identifying patients who may be vulnerable before an event happens.

We also observed more noticeable and statistically significant metabolic changes during and after embolic events, reflecting the widespread systemic effects of pulmonary embolism. These patterns were clearly captured in the longitudinal data. Taken together, our results show that lipid-focused diagnostics and treatments should be a priority in future efforts to prevent pulmonary embolism and improve outcomes for patients with COVID-19.

## Author Contributions

Conceptualization, TH, HE, DG; Data curation, ZZ, LL, MS, WY, CN, JG, LA; Formal analysis, MV, MJ, AK, HE ; Funding acquisition, TH, HE, DG, AH, AK ; Investigation, MV, MJ, AH, AK, HE, DG, TH; Methodology, ; Project administration,; Resources, AH, LL, YR, DG, HE, TH; Software, MV, AK ; Supervision, TH, HE, AK, DG ; Visualization, MV; Writing—original draft, MV, MJ, NK, ; Writing—review & editing, MV, MJ, AK, NK, HE, TH, AH . All authors have read and agreed to the published version of the manuscript.

## Acknowledgments

We gratefully acknowledge the patients whose generous participation and trust made this study possible.

## Institutional Review Board Statement**: -**

### Informed Consent Statement

Informed consent was obtained from all subjects involved in the study.

### Data Availability Statement

All data utilised in the statistical analyses are available as part of the Supplementary files.

## ABBREVIATIONS

-O: alkyl substituent
-P: alkenyl substituent
11,12-DiHETrE: 11,12-dihydroxy-5Z,8Z,14Z-eicosatrienoic acid
13-HODE: Coriolic acid
14,15-DiHETE: (+/-)-14,15-dihydroxy-5Z,8Z,11Z,17Z-eicosatetraenoic acid
15-HETE: 15-hydroxy-5Z,8Z,11Z,13E-eicosatetraenoic acid
15-HETRrE: 15S-hydroxy-8Z,11Z,13E-eicosatrienoic acid
9,10-DiHoME: 9,10-dihydroxy-12Z-octadecenoic acid
9-HOTrE: 9S-hydroxy-10E,12Z,15Z-octadecatrienoic acid
CE: Cholesterol esters
Cer: Ceramides
COVID-19: Coronavirus Disease 2019
DCER: Dihydroceramides
DG: Diacyclglycerols
EDTA: ethylenediaminetetraacetic acid
FA: Free fatty acids
GluCer: Glucoceramides
LacCer: Lactosylceramides
LC-MS: Liquid Chromatography–Mass Spectrometry
LPA: Lysophosphatidic acids
LPC: Lysophosphatidylcholines
LPE: Lysophosphatidylethanolamines
LPG: Lysophosphatidylglycerols
LPI: Lysophosphatidylinositols
LPS: Lysophosphatidylserines
PC: Phosphatidylcholines
PE: Phosphatidylethanolamines
PG: Phosphatidylglycerols
PI: Phosphatidylinositols
PS: Phosphatidylserines
SARS-CoV-2: Severe Acute Respiratory Syndrome Coronavirus 2
serotinin/Trp: serotonin to tryptophan ratio
SM: Sphingomyelins
SP: Sphingosine
TG: Triglycerides
Trp/LNAA: Tryptophan to Large Neutral Amino Acids ratio

